# TAU FILAMENTS WITH THE CHRONIC TRAUMATIC ENCEPHALOPATHY FOLD IN A CASE OF VACUOLAR TAUOPATHY WITH *VCP* MUTATION D395G

**DOI:** 10.1101/2024.04.01.587539

**Authors:** Chao Qi, Ryota Kobayashi, Shinobu Kawakatsu, Fuyuki Kametani, Sjors H.W. Scheres, Michel Goedert, Masato Hasegawa

## Abstract

Dominantly inherited mutation D395G in the gene encoding valosin-containing protein causes vacuolar tauopathy, a type of behavioural-variant frontotemporal dementia, with marked vacuolation and abundant filamentous tau inclusions made of all six brain isoforms. Here we report that tau inclusions were concentrated in layers II/III of the frontotemporal cortex in a case of vacuolar tauopathy. By electron cryo-microscopy, tau filaments had the chronic traumatic encephalopathy (CTE) fold. Tau inclusions of vacuolar tauopathy share this cortical location and the tau fold with CTE, subacute sclerosing panencephalitis and amyotrophic lateral sclerosis/parkinsonism-dementia complex, which are believed to be environmentally induced. Vacuolar tauopathy is the first inherited disease with the CTE tau fold.

## INTRODUCTION

Dominantly inherited mutation D395G in the gene encoding valosin-containing protein (*VCP*) has been described as the cause of an inherited form of behavioural-variant frontotemporal dementia (FTD) in three families from Greece, the US and Japan [5,19,33]. By histology and immunoblotting, abundant neuronal vacuoles and tau protein inclusions made of all six brain tau isoforms were in evidence in the Greek and US families, resulting in the naming of this condition as vacuolar tauopathy [5].

Previously, abundant tau inclusions were described in cases with mutations in genes other than *MAPT*, the tau gene. They include Alzheimer’s disease (AD) (APP and presenilin genes), familial British and Danish dementias (BRI gene), and cases of Gerstmann-Sträussler-Scheinker disease (prion protein gene) [20]. In these diseases, abundant extracellular deposits of various proteins (Aβ, BRI and prion protein) are present alongside intraneuronal tau inclusions. By electron cryo-microscopy (cryo-EM), the Alzheimer tau fold is characteristic of these diseases [8,37].

The Alzheimer tau fold also characterises what has been called primary age-related tauopathy (PART) [41], a sporadic condition where tau inclusions form in an age-related manner, in the absence of extracellular deposits [3]. It follows that the tau inclusions that form in most people as a function of age have the Alzheimer fold.

Mutations in *MAPT* give rise to frontotemporal dementia and parkinsonism linked to chromosome 17 (FTDP-17T), with abundant tau inclusions in brain cells, in the absence of extracellular deposits [10]. So far, cryo-EM has shown the presence of the Pick fold [40] and the argyrophilic grain disease fold [42] in cases of FTDP-17T. Vacuolar tauopathy is the first disease caused by a mutation in a gene other than *MAPT* that results in the formation of abundant neuronal tau inclusions, in the absence of extracellular deposits.

By cryo-EM, tau filaments that are made of all six brain isoforms fall into two groups that consist of the Alzheimer and the chronic traumatic encephalopathy (CTE) folds [37]. The latter is also found in subacute sclerosing panencephalitis (SSPE) [31] and the amyotrophic lateral sclerosis/parkinsonism dementia complex (ALS/PDC) [32]. The CTE tau fold is typical of diseases with abundant inclusions in layers II/III of the neocortex. It consists mostly of repeats three and four, and 10-13 amino acids after repeat four (the longest human brain tau isoform of 441 amino acids has four microtubule-binding repeats of 31 or 32 amino acids each in its C-terminal half) [7].

Here we characterised the neuropathology and the cryo-EM structures of tau filaments from a previously described individual with mutation D395G in *VCP* [19]. A Japanese man died aged 63 after an 18-year history of personality changes and cognitive impairment. Tau inclusions were present in the neocortex, where they were most abundant in layers II/III. Vacuolation was observed mostly in brain regions with few tau inclusions. By cryo-EM, the tau fold was identical to that in CTE, SSPE and ALS/PDC. This is the first inherited tauopathy with the CTE fold.

## MATERIALS AND METHODS

### Clinical presentation

The individual was a man with a heterozygous D395G mutation in *VCP* who developed behavioural changes and cognitive impairment around age 45 [19]. He died of aspiration pneumonia aged 63.

### Immunohistochemistry

Neuropathological examination was carried out as described [16]. Briefly, one brain hemisphere was fixed in 10% neutral buffered formalin and cut into slices of 0.5 cm, whereas the other hemisphere was frozen. Tissue blocks were obtained from approximately thirty regions, including cerebral cortex, basal ganglia, brainstem and cerebellum. The tissues were embedded in paraffin and sectioned at 7 μm for Gallyas-Braak and 4 μm for hematoxylin-eosin (HE) staining and immunolabelling. The following primary antibodies were used: AT8, to detect hyperphosphorylated tau (1:1,000, Innogenetics); anti-Aβ(11-28) (1:400, Immuno-Biological Laboratories); pSyn64, to detect α-synuclein phosphorylated at S129 (1:10,000, Fujifilm); pTDP-43, to detect TDP-43 phosphorylated at S409 and S410 (1:5,000, Cosmo Bio); anti-glial fibrillary acidic protein (1:400, Leica Biosystems); anti-Iba1 (1:1,000, Fujifilm). Primary antibody binding was detected using peroxidase-labelled streptavidin biotin kits (Nichirei Histofine Simole Stain). Diaminobenzidine was used for colour development and the slides were counterstained with HE. Antigen retrieval used pretreatment with formic acid pretreatment [for Aβ (11-28)] or autoclaving [for pSyn64, pTDP-43 and GFAP].

### Filament extraction from brain

Sarkosyl-insoluble material was extracted from the frontal and temporal cortex of the brain of the individual with mutation D395G in *VCP*, as well as from the temporal cortex of a neuropathologically confirmed case of sporadic AD, essentially as described [43]. Briefly, tissues were homogenised with a Polytron in 40 vol (w/v) extraction buffer consisting of 10 mM Tris-HCl, pH 7.4, 0.8 M NaCl, 10% sucrose and 1 mM EGTA. Homogenates were brought to 2% sarkosyl and incubated for 30 min at 37° C. Following a 10 min centrifugation at 27,000 g, the supernatants were spun at 257,000 g for 30 min. Pellets were resuspended in 2 ml extraction buffer containing 1% sarkosyl and centrifuged at 166,000 g for 20 min. The resulting pellets were resuspended in 20 μl buffer containing 20 mM Tris-HCl, pH 7.4, 100 mM NaCl and used for subsequent analysis.

### Immunoblotting

Immunoblotting was carried out as described [43]. Samples were run on 5-20% gradient gels (Fuji Film). Proteins were then transferred to a polyvinylidene difluoride (PVDF) membrane and incubated with phosphorylation-dependent anti-tau mouse monoclonal antibody AT8 (1:1,000) overnight at room temperature. Following washing in PBS, the membranes were incubated with biotinylated anti-mouse antibody (Vector, 1:500) for 2h at room temperature, followed by a 30 min incubation with avidin-biotin complex and colour development using NiCl_2_-enhanced diaminobenzidine.

### Mass spectrometry

Sarkosyl-insoluble fractions from the frontal cortex of the individual with mutation D395G in *VCP* were treated with 70% formic acid for 1h at room temperature, diluted with water and dried. For trypsin digestion, 50 mM triethylammonium bicarbonate and 1 μg of Trypsin/Lys-C mix (Promega) were added. Each mixture was incubated at 37° C for 20 h. Following digestion, 2 μl of 100 mM dithiothreitol were added and the incubation continued at 100° C for 5 min. The samples were dried and stored at-80° C until use. Masss spectrometry was carried out as described [15].

### Electron cryo-microscopy

Cryo-EM grids (Quantifoil 1.2/1.3, 300 mesh) were glow-discharged for 1 min using an Edwards (S150B) sputter coater. Three μl of the sarkosyl-insoluble fractions were applied to the glow-discharged grids, followed by blotting with filter paper and plunge freezing into liquid ethane using a Vitrobot Mark IV (Thermo Fisher Scientific) at 4° C and 100% humidity. Cryo-EM images were acquired on a Titan Krios G2 microscope (Thermo Fisher Scientific) operated at 300 kV and equipped with a Falcon-4i electron detector. Images were recorded for 2s in electron event representation format [11], with a total dose of 40 electrons per A^2^ and a pixel size of 0.824 Å. See Supplementary Table 1 for further details.

### Data processing

Datasets were processed in RELION using standard helical reconstruction [12,18]. Movie frames were gain-corrected, aligned and dose-weighted using RELION’s own motion correction programme [47]. Contrast transfer function (CTF) was estimated using CTFFIND4.1 [34]. Filaments were picked by hand and segments were extracted with a box size of 1,024 pixels, prior to downsizing to 256 pixels. Reference-free 2D classification was carried out and selected class averages were re-extracted using a box size of 400 pixels. Initial models were generated *de novo* from 2D class average images using relion_helix_inimodel2d [36]. Three-dimensional refinements were performed in RELION-4.0 and the helical twist and rise refined using local searches. Bayesian polishing and CTF refinement were used to further improve resolutions [48]. The final maps were sharpened using standard post-processing procedures in RELION-4.0 and resolution estimates were calculated based on the Fourier shell correlation (FSC) between two independently refined half-maps at 0.143 (Supplementary Figure 1) [35]. We used relion_helix_toolbox to impose helical symmetry on the post-processed maps.

### Model building and refinement

Atomic models were built manually using Coot [6], based on published structures (CTE Type I, PDB: 6NWP; CTE Type II, PDB: 6NWQ; CTE Type III, PDB: 8OT9) [7,32]. Model refinements were performed using ISOLDE [4], *Servalcat* [46] and REFMAC5 [25,26]. Models were validated with MolProbity [2]. Figures were prepared with ChimeraX [28] and PyMOL [39].

## RESULTS

At autopsy, the brain from the individual with mutation D395G in *VCP* weighed 968 g. A side view of the intact brain showed atrophy of the frontal cortex (Figure 1a). Coronal sections revealed moderate atrophy of the frontal cortex, with mild atrophy of temporal and parietal cortex, and no atrophy of occipital cortex, hippocampus, amygdala or basal ganglia (Figure 1b-d). There was depigmentation of the locus coeruleus, but not the substantia nigra.

**Figure 1.**
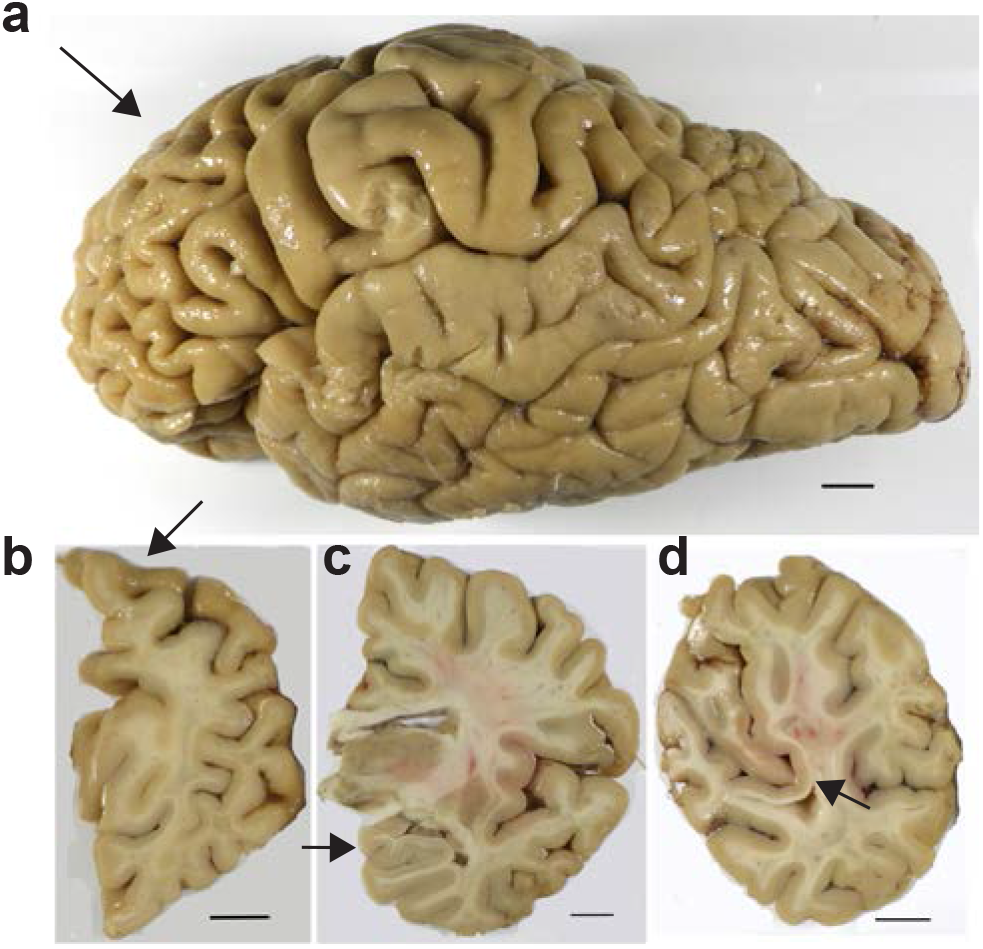
Formalin-fixed brain from the individual with mutation D395G in VCP. a, Side view of the brain, showing atrophy of the frontal cortex (arrowed). b, Coronal section of the cerebral hemispheres with atrophy of the dorsal portion of the frontal lobe (arrowed). c, Coronal section at the level of the thalamus, indicating preservation of the hippocampus (arrowed). d, Coronal section showing preservation of the occipital cortex, including the primary visual cortex (arrowed). Scale bars, 10 mm.

Abundant tau-immunoreactive, Gallyas-Braak silver-positive neurofibrillary lesions, including ghost tangles, were observed in frontal (Figure 2a,d; Figure 3), temporal (Figure 2b,e; Figure 4) and parietal (Figure 2c,f) cortex. They were rare in occipital cortex and hippocampus (Figure 2b,c,e,f; Figure 5b,c,e,f,h,i). Neuronal cell loss (Figure 3a,f,k; Figure 4a,f,k) and neurofibrillary lesions were concentrated in the upper cortical layers of frontal (Figure 3b,c,g,h) and temporal (Figure 4b,c,g,h) cortex. They were also abundant in nucleus basalis of Meynert, thalamus, substantia nigra, caudate nucleus, locus coeruleus, pons and medulla oblongata. The cerebellum was well preserved, with only a few neurofibrillary lesions in the dentate nucleus.

**Figure 2.**
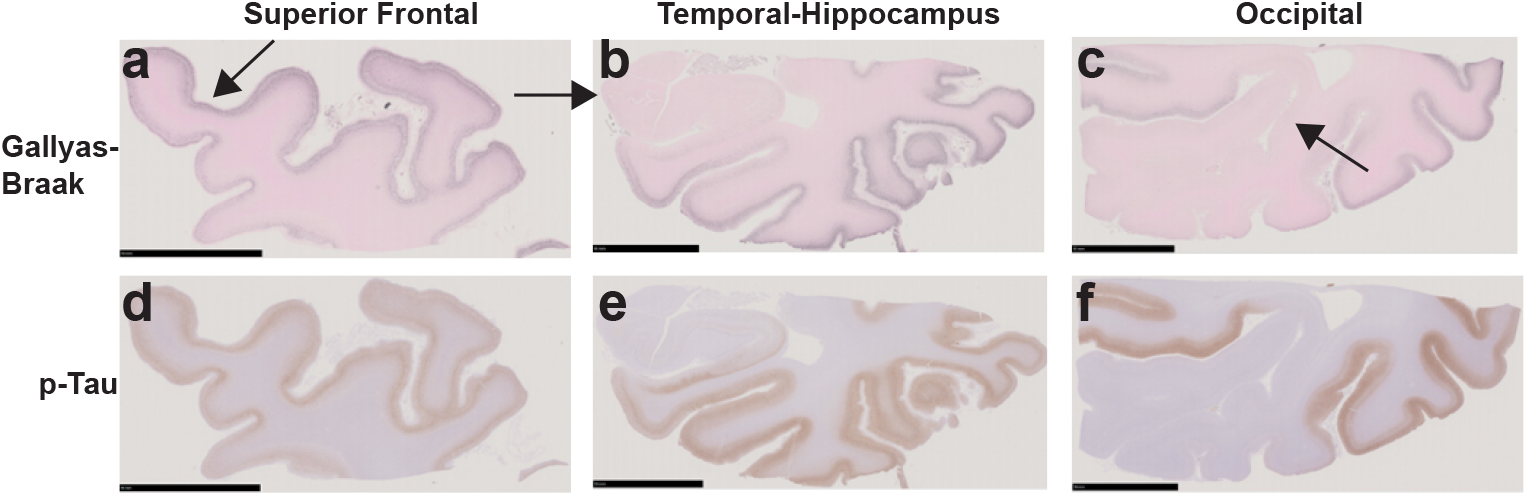
Gallyas-Braak silver and pTau (AT8) staining of coronal brain sections of the superior frontal, the temporal-hippocampus and the occipital cortex from the individual with mutation D395G in VCP. a-c, Gallyas-Braak silver shows numerous lesions in the superficial layers of the atrophic superior frontal cortex (arrowed) (d), inferior to middle temporal (e) and occipital (f) cortex, whereas hippocampus (arrowed) and primary visual cortex (arrowed) are intact. Scale bars, 10 mm. d-f, Similar to Gallyas-Braak silver, AT8 immunostaining shows strong staining of frontal, temporal and parietal cortical layers, with sparing of the hippocampus and the primary visual cortex. Scale bars, 10 mm.

**Figure 3.**
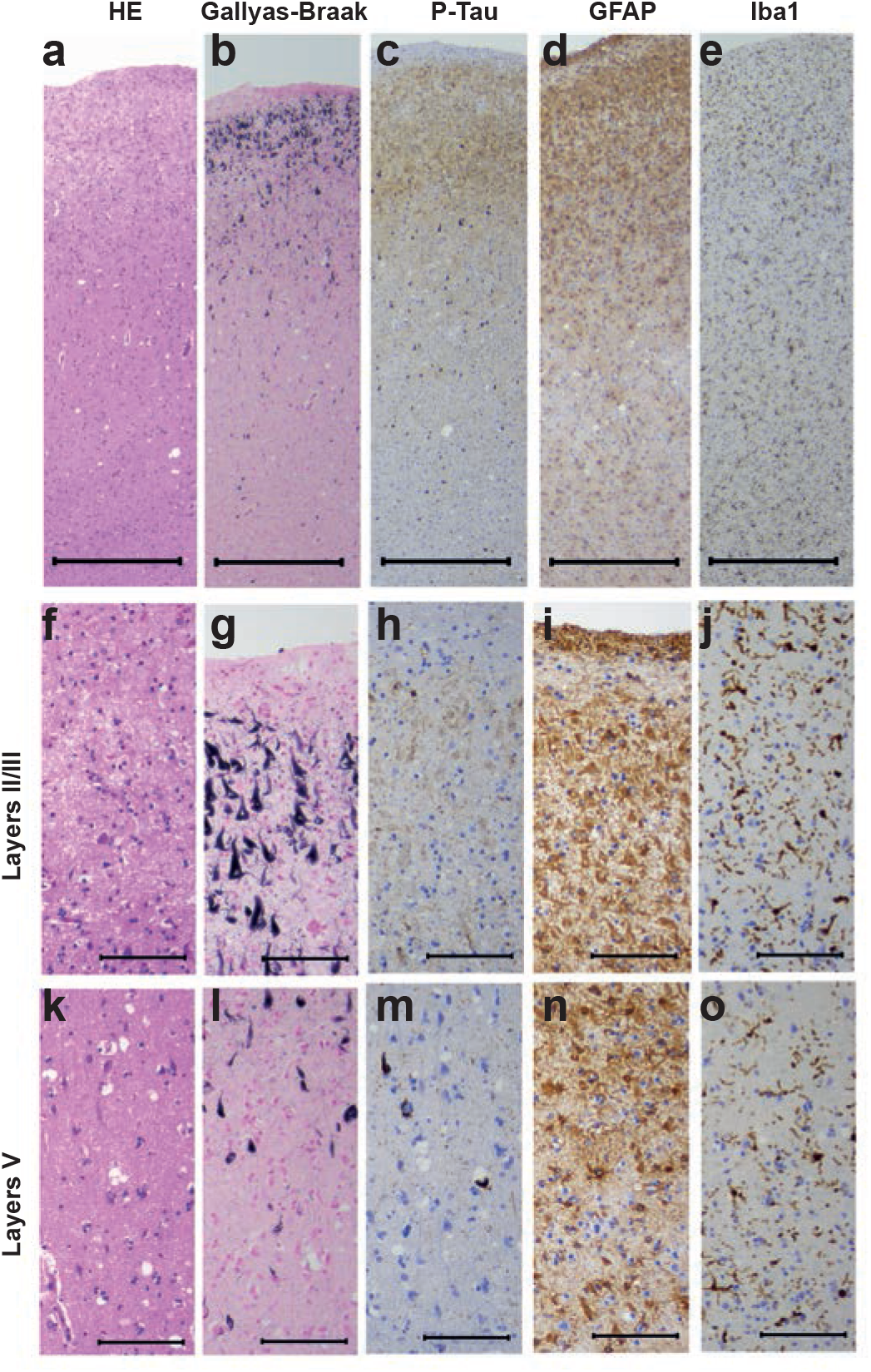
Staining of the superior frontal cortex in vacuolar tauopathy. Nerve cell loss and gliosis are in seen in layers II/III (a,d f,i), where abundant tau-immunoreactive neurofibrillary lesions are in evidence (b,c,g,h). Fewer neurofibrillary lesions are seen in layer V (l,m). Mild vacuolar changes are present in the deep cortical layers (k). Astrogliosis and microglial changes are most severe in the superficial cortical layers (d,e,i j,n,o). HE staining (a,f,k); Gallyas-Braak silver (b,g,l); pTau (AT8) (c,h,m); GFAP (d,i,n); Iba1 (e,j,o). Scale bars: 500 μm (a-e), 100 μm (f-o).

**Figure 4.**
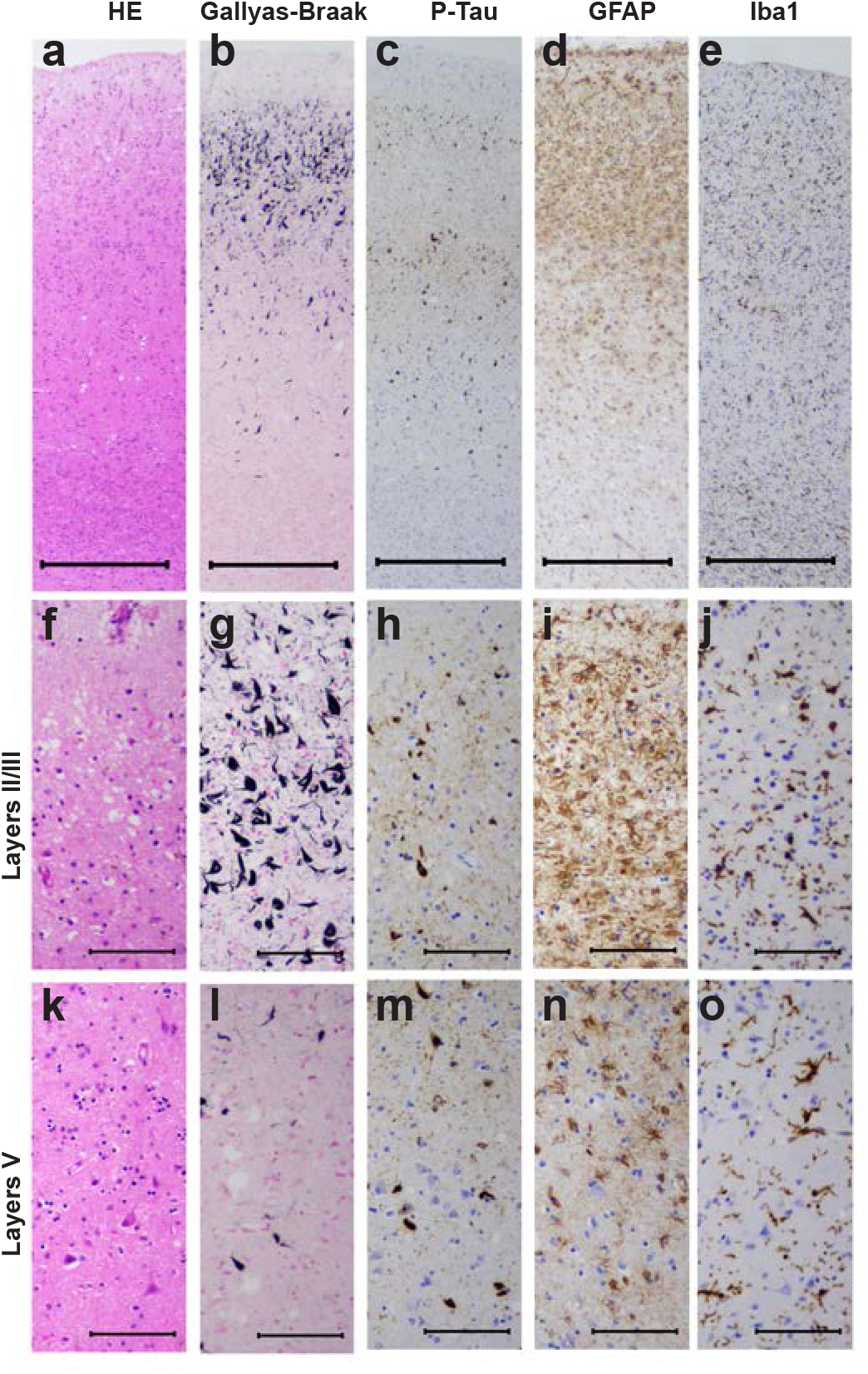
Staining of the middle temporal cortex in vacuolar tauopathy. Nerve cell loss and gliosis are seen in layers II/III (a,d,f,i), where abundant tau-immunoreactive neurofibrillary lesions are in evidence (b,c,g,h). Fewer neurofibrillary lesions are seen in layer V (l,m). Mild vacuolar changes are present in the superficial cortical layers (f). Astrogliosis and microglial changes are most severe in the superficial layers of the temporal cortex (d,e,i,j,n,o). HE staining (a,f,k); Gallyas-Braak silver (b,g,l); pTau (AT8) (c,h,m); GFAP (d,i,n); Iba1 (e,j,o). Scale bars: 500 μm (a-e), 100 μm (f-o).

**Figure 5.**
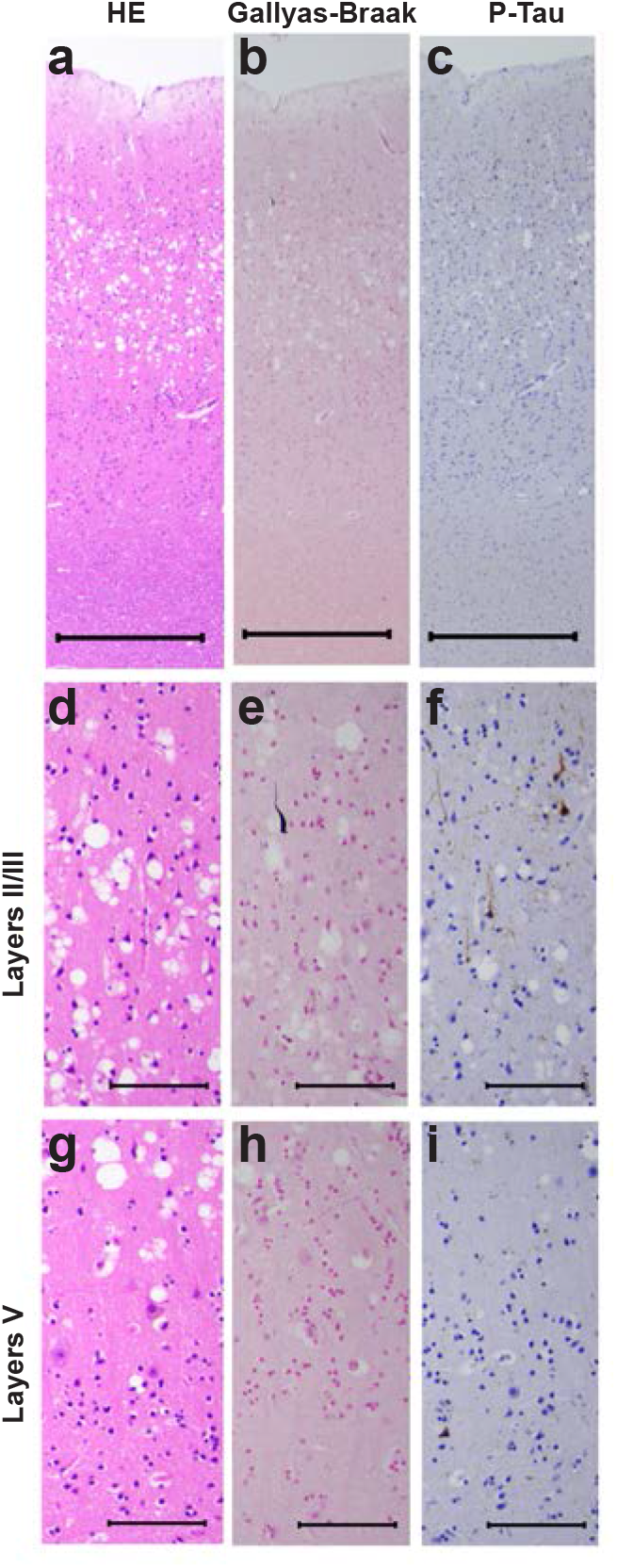
Staining of the primary visual cortex in vacuolar tauopathy. Neurofibrillary lesions are almost absent (b,c,e,f,h,i). Severe vacuolar changes are present, in particular in the superficial cortical layers (a,d,g). HE staining (a,d,g); Gallyas-Braak silver (b,e,h); pTau (AT8) (c,f,i). Scale bars: 500 μm (a-c), 100 μm (d-i).

Severe vacuolar changes were observed predominantly in the superficial layers of the primary visual cortex (Figure 5a,d), where neurofibrillary lesions (Figure 5b,c,e,f,h,i) were almost absent. Mild vacuolar changes were also present in frontal (Figure 3k) and temporal (Figure 4f) cortex. Astrogliosis and activation of microglia were evident in the frontal (Figure 3d,e,i,j,n,o) and temporal (Figure 4d,e,i,j,n,o) cortex. This case is thus another example of vacuolar tauopathy caused by mutation D395G in *VCP* .

Immunoblotting of the sarkosyl-insoluble fraction from the temporal cortex of the individual with vacuolar tauopathy with anti-tau antibody AT8 showed the presence of strong bands of 60, 64 and 68 kDa and a weak band of 72 kDa. This pattern was identical to that from a case of sporadic AD (Figure 6). The post-translational modifications of sarkosyl-insoluble tau extracted from the frontal cortex of the individual with vacuolar tauopathy (Supplementary Table 2) were also similar to those reported in AD [15,45]. No Aβ, α-synuclein or TDP-43 inclusions were observed.

**Figure 6.**
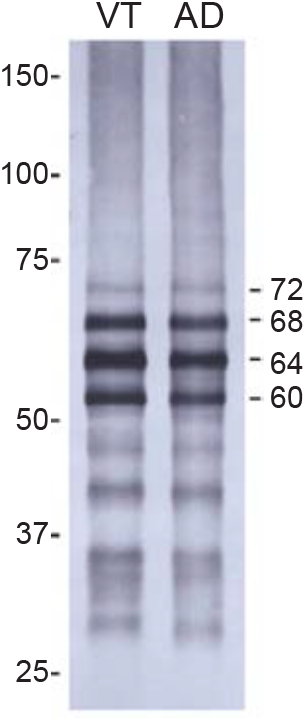
Immunoblotting of sarkosyl-insoluble fractions from the temporal cortex of the individual with vacuolar tauopathy (VT) and a case of sporadic Alzheimer’s disease (AD). Phosphorylation-dependent anti-tau antibody AT8 was used. Note the presence of strong bands of 60, 64 and 68 kDa and a weaker band of 72 kDa (indicated on the right).

We used cryo-EM to determine the structures of tau filaments extracted from the frontal and temporal cortex of the individual with vacuolar tauopathy. Three Types of filaments were observed that were made of two identical protofilaments arranged in different ways (Figure 7a,b). All three filament types were found in the frontal cortex, with Type III being absent from the temporal cortex. Filament structures were determined to resolutions ranging from 2.3 to 3.4 Å, which allowed us to establish their identity as CTE filament Types I-III [7,32] (Figure 7c). The ordered cores of the filaments span residues K274-R379 of three-repeat tau and residues S305-R379 of four-repeat tau. The root mean square deviation (rmsd) between Cα atoms of Type I and CTE Type I filaments filaments was 0.278 Å; that between Type II and CTE Type II filaments filaments was 0.566 Å and that between Type III and CTE Type III filaments was 0.568 Å.

**Figure 7.**
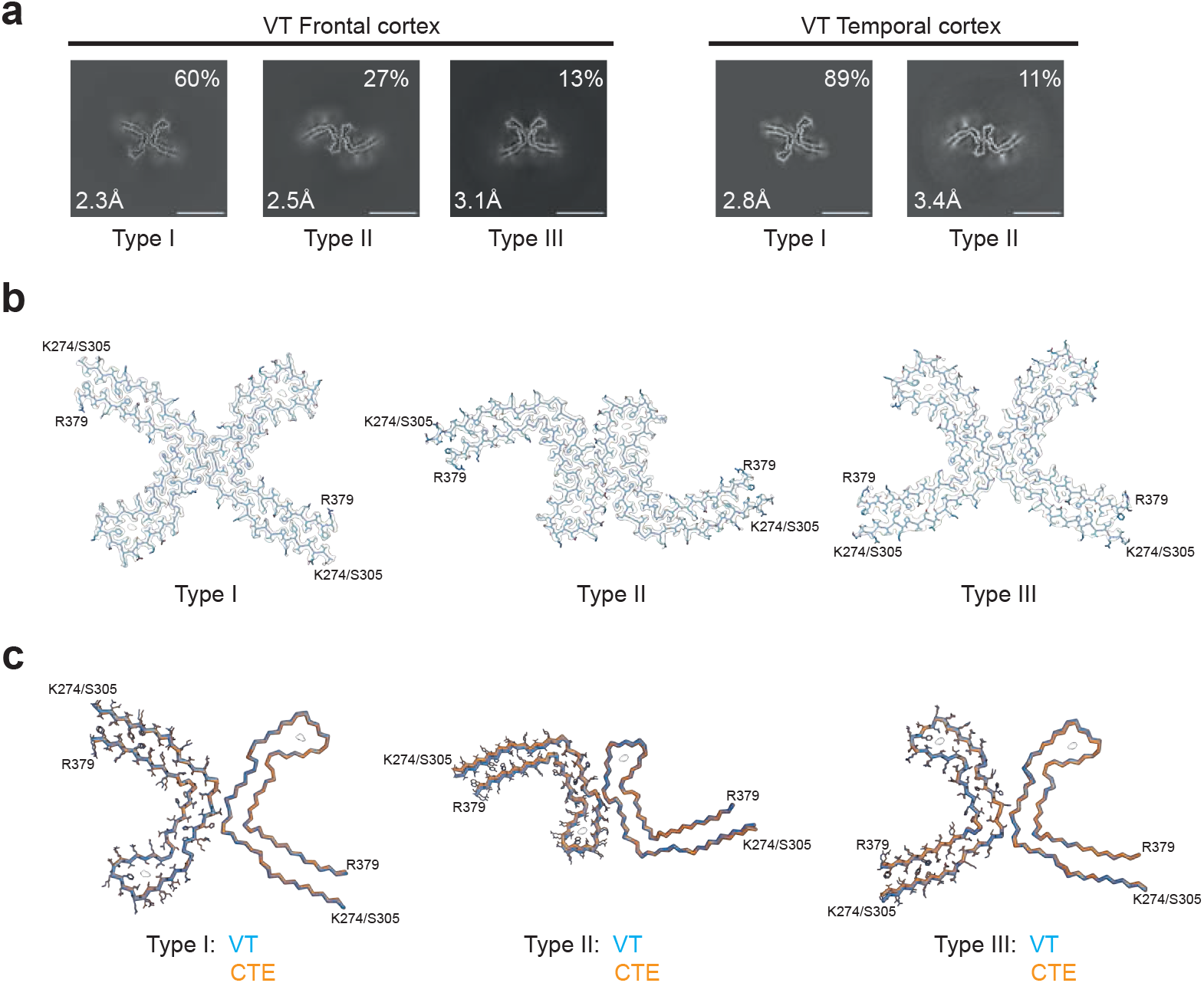
Cryo-EM cross-sections and structures of tau filaments from vacuolar tauopathy. a, Cross-sections through the cryo-EM reconstructions, perpendicular to the helical axis and with a projected thickness of approximately one rung, are shown for frontal and temporal cortex. Three filament types were present (Type III was only found in the frontal cortex). They are made of two identical protofilaments that are arranged in different ways. Resolutions (in Å) and percentages of filament types are indicated in the bottom left and top right, respectively. Scale bar, 10 nm. b, Cryo-EM density maps and models of Type I, Type II and Type III tau filaments from the case with vacuolar tauopathy. c, Type I, Type II and Type III filaments from the case with vacuolar tauopathy (in blue) overlaid with CTE Type I, Type II and Type III filaments from a case with CTE (in orange). The ordered cores of the filaments extend from tau K274/S305-R379.

## DISCUSSION

Mutation D395G in *VCP* causes vacuolar tauopathy, a type of frontotemporal dementia with widely distributed and abundant filamentous tau inclusions [5]. Here we report the neuropathology from a previously described case with mutation D395G in *VCP* [19] and establish that it is a case of vacuolar tauopathy.

In the neocortex, Gallyas-Braak silver-positive tau inclusions were concentrated in the superficial layers. Inclusions were present in frontal, temporal and parietal cortex, as well as in several other brain regions. The frontal cortex was moderately atrophic. Immunoblotting showed a pattern of tau bands like that in AD and CTE, consistent with the presence of all six brain tau isoforms [9,38]. By mass spectrometry, the post-translational modifications of sarkosyl-insoluble tau extracted from the frontal cortex of the individual with vacuolar tauopathy were like those reported in AD [15,45].

Neuronal vacuolisation and neurofibrillary degeneration were inversely related. Thus, abundant vacuoles were found mostly in regions that had only few neurofibrillary lesions, such as the occipital cortex. Conversely, regions with abundant tau inclusions, such as the frontal and the temporal cortex, had only few vacuoles. This suggests that the D395G mutation in *VCP* causes neuronal vacuolisation and neurofibrillary degeneration through distinct mechanisms. Astrocytosis and microgliosis were seen in regions with neurofibrillary degeneration, as is the case of other diseases with neurofibrillary tau pathology [17].

These findings are reminiscent of those reported previously in four individuals from two families with the *VCP* mutation D395G [5]. Vacuolisation appeared to be endocytic in origin and was found mostly in cells that were not destined for neurodegeneration. The previous work also hypothesised that VCP might be a disaggregase for assembled and ubiquitinated tau, with assembled tau accumulating as the result of a partial loss-of-function resulting from mutation D395G. However, it was surprising that this mutation did not appear to affect other proteins known to aggregate in a polyubiquitinated form in the human brain, such as α-synuclein and TDP-43. *In vitro* experiments have shown that VCP prevents the seeding, not only of assembled tau [5], but also of assembled α-synuclein and TDP-43 [49].

VCP is an AAA^+^ ATPase that unfolds ubiquitinated proteins [1]. Besides mutation D395G, other mutations in *VCP* cause multisystem proteinopathy, a degenerative disease affecting muscle and bone, that can also present as frontotemporal dementia with TDP-43 inclusions [29,44]. Unlike D395G, which results in a reduction in the ATPase activity of VCP, these mutations increase the ATPase activity and are believed to result in a gain-of-toxic function of VCP [5].

We used cryo-EM to determine the fold of tau filaments extracted from the frontal and temporal cortex of the individual with vacuolar tauopathy. The filaments had the CTE tau fold [7], which we also identified in cases of SSPE [31] and ALS-PDC [32]. Types I-III of CTE filaments [7,31] were present in the frontal cortex, with only Types I and II being found in the temporal cortex. They are molecular polymorphs that consist of two identical protofilaments linked in different ways. In these diseases, more filamentous tau inclusions are found in layers II/III of the cerebral cortex than in layer V [13,23,24]. This is unlike AD, where tau inclusions are more abundant in layer V [27].

So far, there has been an absolute correlation between the cortical localisation of tau inclusions and the presence of the CTE or the Alzheimer fold. It follows that the CTE tau fold may also form in other diseases with a predominance of tau inclusions in cortical layers II/III, such as Postencephalitic Parkinsonism [14] and the Nodding Syndrome [30]. However, unlike CTE, SSPE and ALS/PDC, which are believed to have mainly environmental causes, vacuolar tauopathy is dominantly inherited. This is the first inherited condition with the CTE tau fold. It remains to be determined if tau filaments extracted from the brains of other cases with vacuolar tauopathy [5] also carry the CTE fold.

The CTE tau fold differs from the Alzheimer fold by having a more open conformation of the β-helix region, which contains an internal density of unknown identity [7]. In the presence of NaCl, recombinant tau comprising residues 297-391 assembles into filaments with the CTE fold, but in the presence of MgCl_2_, the Alzheimer fold forms [21]. Both folds assemble from a shared transient first intermediate amyloid filament, followed by multiple different polymorphic filamentous intermediates [22].

It remains to be determined how mutation D395G in *VCP* leads to the presence of tau filaments with the CTE fold. VCP may be a disaggregase of this fold. On the other hand, we have previously hypothesised that the CTE tau fold could form in response to separate insults, which might be linked by specific neuroinflammatory changes that differ from those common to all tauopathies [32]. A partial loss-of-function of VCP could therefore give rise to mechanisms that result in the formation of the CTE fold without requiring a direct interaction with tau.

## Acknowledgements

This work was supported by the Electron Microscopy Facility of the MRC Laboratory of Molecular Biology. We thank Jake Grimmett, Toby Darling and Ivan Clayson for help with high-performance computing, and Takumi Kitaoka and Mitsuru Futakuchi (Yamagata University School of Medicine) for help with neuropathology. We also thank Hiroya Naruse and Tatsushi Toda (University of Tokyo) for genomic analysis. For open access, the MRC Laboratory of Molecular Biology has applied a CC BY public copyright licence to any Author Accepted Manuscript version arising.

## Author contributions

RK and SK identified the patient and performed genetic analysis and neuropathology; MH prepared filaments and performed immunoblots; CQ performed cryo-EM data acquisition and structure determination; SHWS, MG and MH supervised the project and all authors contributed to the writing of the manuscript.

## Declarations

### Ethics approval and consent

Studies carried out at Yamagata University were approved through the Institution’s ethical review process. Informed consent was obtained from the patient’s next of kin. This study was approved by the Cambridgeshire 2 Research Ethics Committee (09/H0308/163).

### Open Access

This article is licensed under a Creative Commons Attribution 4.0 International License, which permits use, sharing, adaptation, distribution and reproduction in any medium or format, as long as you give appropriate credit to the original author(s) and the source, provide a link to the Creative Commons licence, and indicate if changes were made. The images or other third party material in this article are included in the article’s Creative Commons licence, unless indicated otherwise in a credit line to the material. If material is not included in the article’s Creative Commons licence and your intended use is not permitted by statutory regulation or exceeds the permitted use, you will need to obtain permission directly from the copyright holder. To view a copy of this licence, visit http://creativecommons.org/licenses/by/4.0/.

---

**Figure S1.**
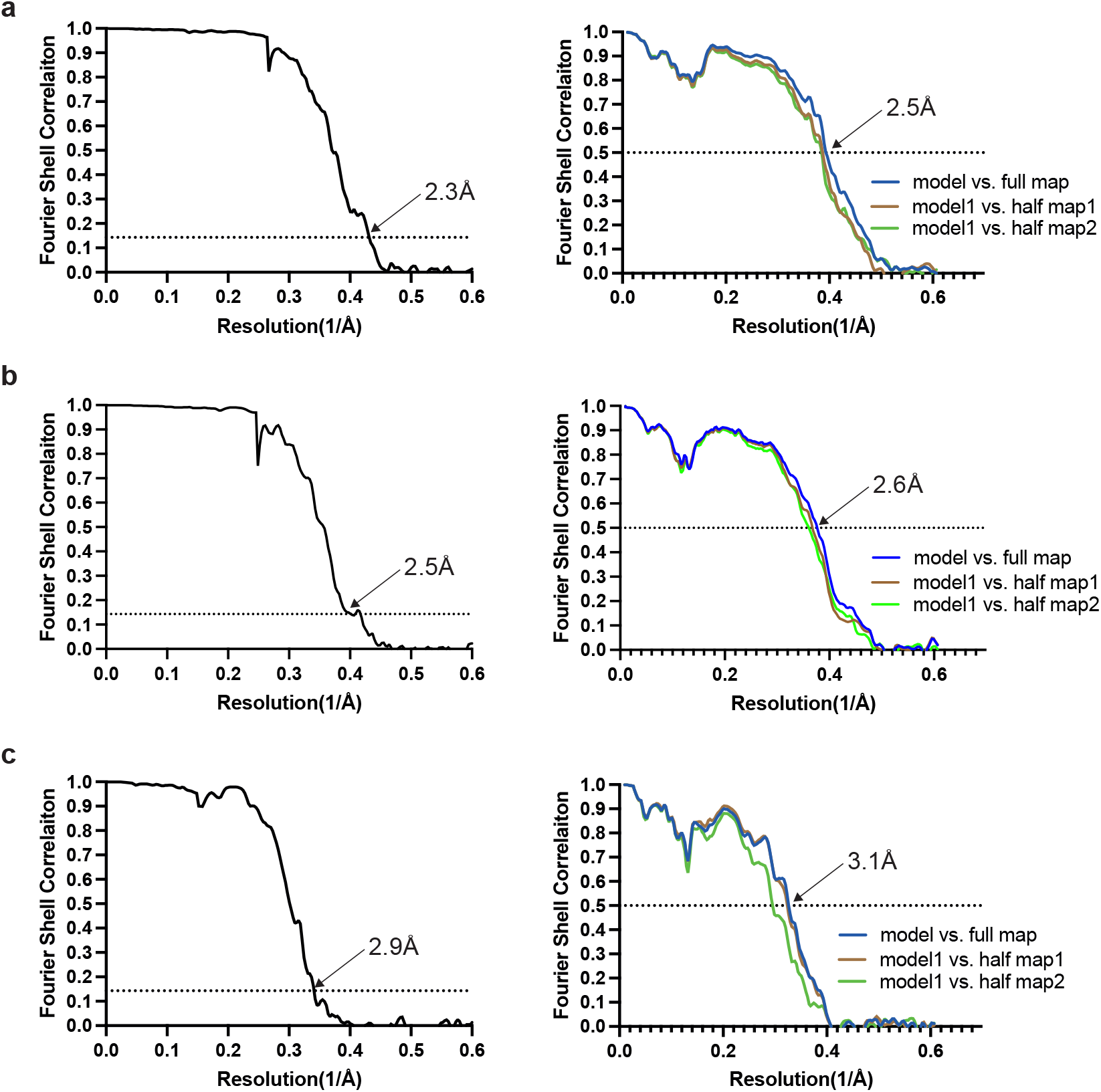
Fourier shell correlation (FSC) curves. FSC curves of cryo-EM maps (left panel) and model to map validation (right panel). a, CTE Type I tau filament from VT; CTE Type II tau filament from VT; CTE Type III tau filament from VT.

**Supplementary Table 1:** Cryo-EM data collection, refinement and validation statistics.

|  |  |  |  |
| --- | --- | --- | --- |
|  | VT Frontal cortex |  |  |
| **Data collection** |  |  |  |
| Microscope | Titan Krios |  |  |
| Voltage (kV) | 300 |  |  |
| Detector | Falcon4 |  |  |
| Magnification | 96,000 |  |  |
| Electron exposure (e-/Å<sup>2</sup>) | 40 |  |  |
| Defocus range (μm) | -1.0 to -2.0 |  |  |
| Pixel size (Å) | 0.824 |  |  |
| **Data processing** | CTE Type I | CTE Type II | CTE Type III |
| Box size (pixel) | 400 | 400 | 400 |
| Symmetry imposed | C1 | C1 | C1 |
| Initial particle images (no.) | 861,191 |  |  |
| Final particle images (no.) | 79,536 | 206,370 | 104,579 |
| Map resolution (Å)<br>FSC threshold 0.143 | 2.3 | 2.5 | 2.9 |
| Helical rise (Å) | 2.4 | 2.4 | 4.8 |
| Helical twist (°) | 179.4 | 179.4 | -1.19 |
| **Refinement** |  |  |  |
| Model resolution (Å)<br>FSC threshold 0.5 | 3.1 | 2.5 | 3.1 |
| Map sharpening *B* factor (Å<sup>2</sup>) | -41 | -47 | -58 |
| Model composition |  |  |  |
| Non-hydrogen atoms | 2870 | 3444 | 5775 |
| Protein residues | 375 | 450 | 750 |
| Ligands | 0 | 0 | 0 |
| *B* factors (Å<sup>2</sup>) |  |  |  |
| Protein | 157.7 | 200.1 | 201.8 |
| R.m.s. deviations |  |  |  |
| Bond lengths (Å) | 0.0069 | 0.0063 | 0.0081 |
| Bond angles (°) | 1.313 | 1.269 | 1.477 |
| Validation |  |  |  |
| MolProbity score | 1.41 | 1 | 2.08 |
| Clashscore | 0.85 | 0 | 3.65 |
| Poor rotamers (%) | 3.03 | 3.03 | 5.26 |
| Ramachandran plot |  |  |  |
| Favored (%) | 95.89 | 97.26 | 94.52 |
| Allowed (%) | 4.11 | 2.74 | 5.26 |
| Disallowed (%) | 0 | 0 | 0 |
| EMDB | 9ERM | 9ERN | 9ERO |
| PDB | EMD-19926 | EMD-19927 | EMD-19928 |

**Supplementary Table 1:**
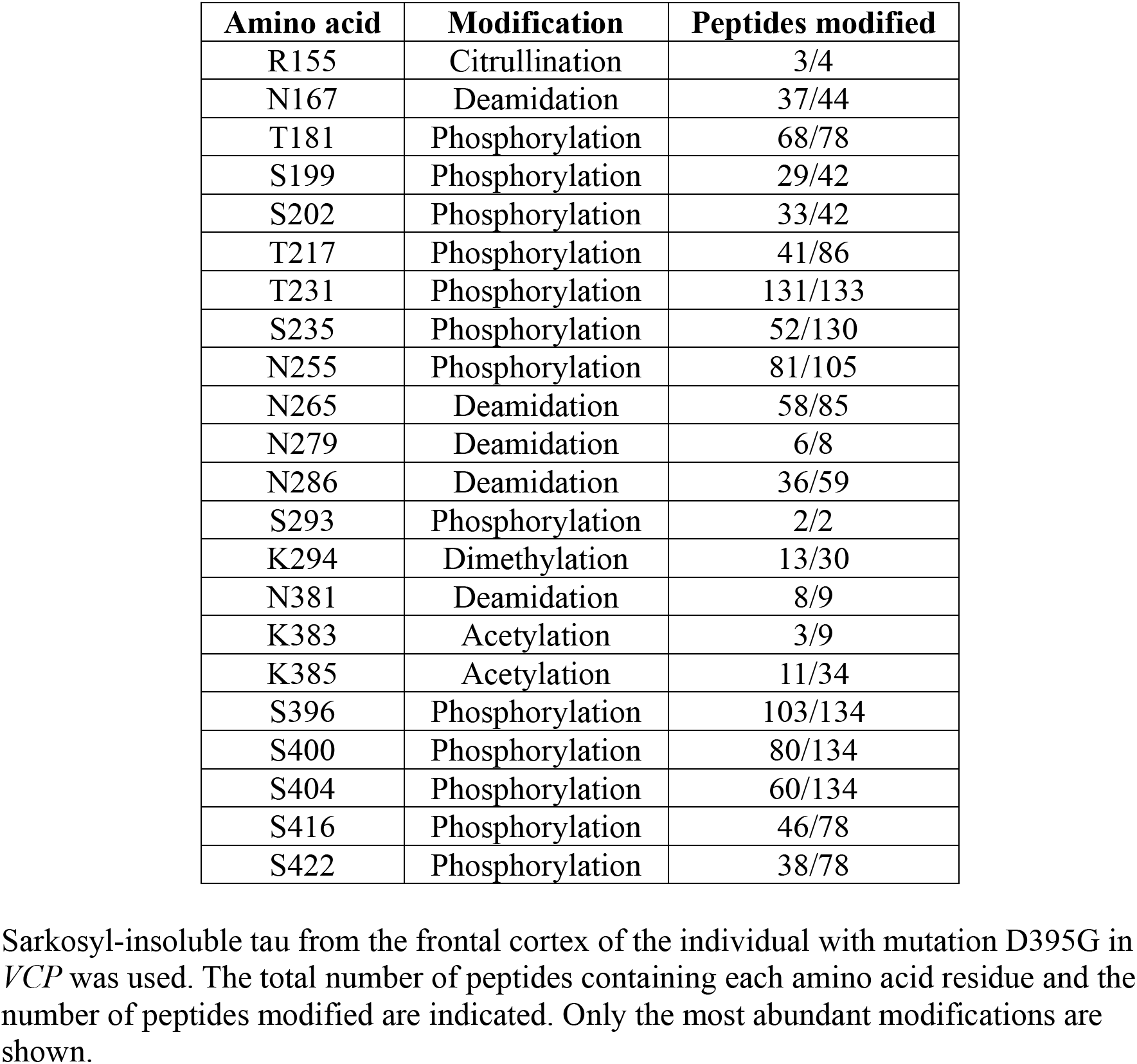
Post-translational modifications.

